# Oncogenic effects of germline mutations in lysosomal storage disease genes

**DOI:** 10.1101/380121

**Authors:** Junghoon Shin, Daeyoon Kim, Hyung-Lae Kim, Murim Choi, Jan O. Korbel, Sung-Soo Yoon, Youngil Koh, on behalf of the PCAWG Germline Cancer Genome Working Group and the ICGC/TCGA Pan-Cancer Analysis of Whole Genomes Network

**Affiliations:** Division of Hematology and Medical Oncology, Department of Internal Medicine, Seoul National University Hospital, Seoul, Korea.; Cancer Research Institute, Seoul National University College of Medicine, Seoul, Korea.; Department of Biochemistry, Ewha Womans University School of Medicine, Seoul, Korea.; Department of Biomedical Sciences, Seoul National University College of Medicine, Seoul, Korea.; European Molecular Biology Laboratory, Genome Biology Unit, 69117, Heidelberg, Germany.; Biomedical Research Institute, Seoul National University College of Medicine, Seoul, Korea.

## Abstract

Clinical observations have indicated that patients with Gaucher disease or Fabry disease are at increased risk of cancer. However, a systematic evaluation of the oncogenic effects of causal mutations of lysosomal storage diseases (LSDs) has been lacking. Here we report a comprehensive association analysis between potentially pathogenic germline mutations in LSD genes and cancer interrogating genomic (or exomic) variant datasets derived from the Pan-Cancer Analysis of Whole Genomes project (case cohort), the 1000 Genomes project (primary control cohort), and the Exome Aggregation Consortium that does not include The Cancer Genome Atlas subset (validation control cohort). We show that potentially pathogenic variants (PPVs) in 42 LSD genes are significantly enriched in cancer patients in a histology-dependent manner, cancer risk is higher in individuals with a greater number of PPVs, and cancer develops earlier in PPV carriers. Analysis of tumor genomic and transcriptomic data from the pancreatic adenocarcinoma cohort revealed potential mechanisms that might be involved in the oncogenic contribution of PPVs. Our findings extend the mechanistic understanding of inherited cancer susceptibility and highlight the promise of harnessing available therapeutic strategies to restore lysosomal function for personalized cancer prevention.

## Introduction

Lysosomal storage diseases (LSDs) comprise more than 50 disorders caused by inborn errors of metabolism, which involve the impaired function of endosome-lysosome proteins.^1^ In LSDs, defects in genes encoding lysosomal hydrolases, transporters, and enzymatic activators result in macromolecule accumulation in the late endocytic system.^2^ The disruption of lysosomal homeostasis is linked to increased endoplasmic reticulum and oxidative stress, which not only is a common mediator of apoptosis in LSDs but also can induce oncogenic cellular phenotype and promote the development of malignancy.^3, 4^

Typical LSD patients have severely impaired organ functions and short life expectancy. However, a considerable number of undiagnosed LSD patients have mildly impaired lysosomal function and survive into adulthood.^1^ These patients are often diagnosed after they develop secondary diseases such as Parkinsonism that is attributable to insidious LSDs.^5^ Clinical observations have shown that patients with Gaucher disease or Fabry disease are at increased risk of cancer,^6, 7^ indicating that dysregulated lysosomal metabolism may contribute to carcinogenesis. However, the precise relationship between lysosomal dysfunction and cancer remains unclear; this uncertainty can be attributed in part to the diverse and nonspecific phenotypes of LSDs and the resulting difficulty in recognizing patients with mild symptoms. The extensive allelic heterogeneity and the complex genotype-phenotype relationships make the diagnosis more challenging.^8^ Furthermore, growing evidence suggests that single allelic loss is functionally significant, even though the impact may not be sufficient to develop overt disease.^9^ Considering the above along with the recessive inheritance nature of most LSDs, we hypothesized that there would be a large number of undetected carriers of causal mutations of LSDs with mild functional impairment, and these carriers would be at increased risk of cancer.

Here we report the results of a comprehensive association analysis between germline mutations in LSD-related genes and cancer using data from global sequencing projects. We show that carriers of potentially pathogenic variants (PPVs) in 42 LSD genes are at increased risk of cancer, cancer risk is higher in individuals with a greater number of PPVs, cancer develops earlier in PPV carriers, and transcriptional misregulation of cancer-promoting signaling pathways might underlie the oncogenic contribution of PPVs. We aimed to elucidate the PPV-cancer association in a histology-specific manner. Potential carcinogenic mechanisms were investigated using tumor genomic and transcriptomic data with a focus on the pancreatic adenocarcinoma.

## Results

### Characteristics of study cohorts

We used matched tumor-normal pair whole genome and tumor whole transcriptome sequence data and clinical and histological annotation of 2,567 cancer patients (Pan-Cancer cohort) from the International Cancer Genome Consortium (ICGC) /The Cancer Genome Atlas (TCGA) Pan-Cancer Analysis of Whole Genomes (PCAWG) project.^10^ As controls, we used publicly available variant call sets from two global sequencing projects of individuals without known cancer histories. The first control dataset comprised 2,504 genomes from the 1000 Genomes project phase 3 (1000 Genomes cohort).^11^ The second dataset included exomes of 53,105 unrelated individuals from a subset of the Exome Aggregation Consortium release 1.0 that did not include TCGA subset (ExAC cohort).^12^

The Pan-Cancer cohort consisted of four populations and 38 histological types of pediatric or adult cancer (Figs. 1a and 1c and Supplementary Table 1). The median age at diagnosis was 60 years (range, 1 to 90). A majority of the patients were Europeans or Americans in most cancer types. The 1000 Genomes cohort comprised five populations (Fig. 1b);^11^ we combined the European and American populations for comparison with the Pan-Cancer cohort. The ExAC cohort included seven populations, among which the Americans and Non-Finnish Europeans together accounted for more than 60% of the entire cohort.^12^

**Fig. 1.**
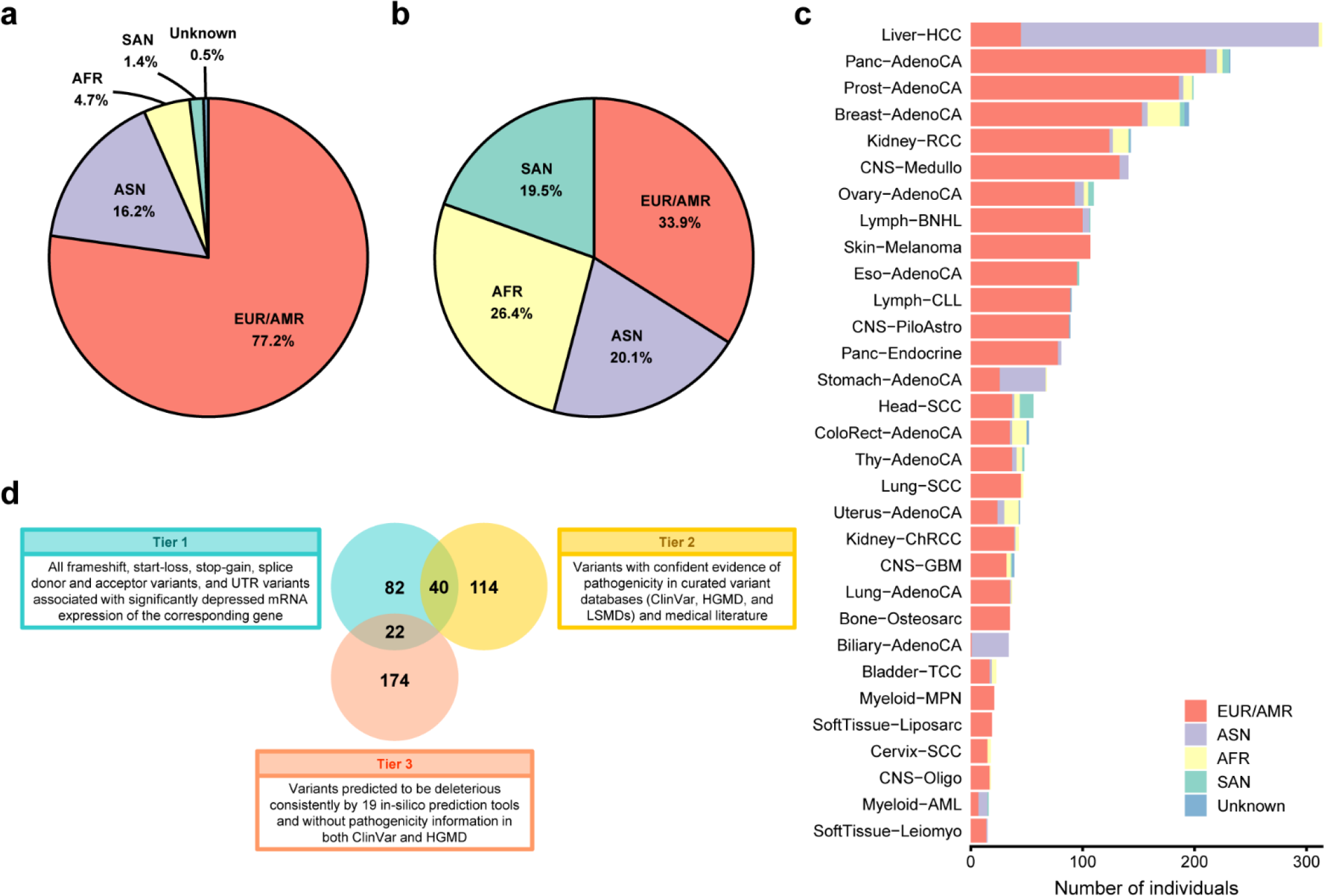
Populations of the Pan-Cancer and 1000 Genomes cohorts and PPV selection criteria. **a**,**b**, Populations comprising the Pan-Cancer cohort (**a**) and the 1000 Genomes cohort (**b**). **c**, Populations comprising each cancer type of the Pan-Cancer cohort (see Supplementary Table 1 for abbreviations of histological subgroups). EUR, European; AMR, American; ASN, East Asian; AFR, African; SAN, South Asian. **d**, Venn diagram of PPVs identified in the Pan-Cancer and 1000 Genomes cohorts grouped into three tiers.

### PPV prevalence in the Pan-Cancer and 1000 Genomes cohorts

Through an extensive literature review, we identified 42 LSD genes (Table 1).^1,8,13–15^ Based on the GRCh37/hg19 genomic coordinates, 7,187 germline single nucleotide variants (SNVs) and small insertions and deletions (indels) were identified in protein-coding regions, essential splice junctions, and 5’ and 3’ untranslated regions (UTRs) in the aggregate variant call set of the Pan-Cancer and 1000 Genomes cohorts (Supplementary Fig. 1). Of those, 4,019 (55.9%) were singletons (variants found in only one individual), and 3′ UTR variants accounted for the largest proportion (37.7%).

**Table 1.** Lysosomal storage disease genes included in this study.

| HGNC Symbol | Chromosome | Associated Lysosomal Storage Disease | Inheritance* |
| --- | --- | --- | --- |
| *AGA* | 4 | Aspartylglycosaminuria | Autosomal recessive |
| *ARSA* | 22 | Metachromatic leukodystrophy | Autosomal recessive |
| *ARSB* | 5 | Mucopolysaccharidosis VI (Maroteaux–Lamy syndrome) | Autosomal recessive |
| *ASAH1* | 8 | Farber lipogranulomatosis | Autosomal recessive |
| *CLN3* | 16 | Neuronal ceroid lipofuscinosis (NCL) 3 (juvenile NCL or Batten disease) | Autosomal recessive |
| *CTNS* | 17 | Cystinosis | Autosomal recessive |
| *CTSA* | 20 | Galactosialidosis | Autosomal recessive |
| *CTSK* | 1 | Pycnodysostosis | Autosomal recessive |
| *FUCA1* | 1 | Fucosidosis | Autosomal recessive |
| *GAA* | 17 | Glycogen storage disease type II (Pompe disease) | Autosomal recessive |
| *GALC* | 14 | Globoid cell leukodystrophy (Krabbe disease) | Autosomal recessive |
| *GALNS* | 16 | Mucopolysaccharidosis IVA (Morquio A syndrome) | Autosomal recessive |
| *GBA* | 1 | Gaucher disease | Autosomal recessive |
| *GLA* | X | Fabry disease | X-linked recessive |
| *GLB1* | 3 | Mucopolysaccharidosis IVB (GM1 gangliosidosis and Morquio B syndrome) | Autosomal recessive |
| *GM2A* | 5 | GM2-gangliosidosis type AB | Autosomal recessive |
| *GNPTAB* | 12 | Mucopolipidosis II (I-cell disease)<br>Mucopolipidosis IIIA (pseudo-Hurler polydystrophy) | Autosomal recessive |
| *GNPTG* | 16 | Mucopolipidosis IIIC (mucopolipidosis III gamma) | Autosomal recessive |
| *GNS* | 12 | Mucopolysaccharidosis IIID (Sanfilippo syndrome D) | Autosomal recessive |
| *GUSB* | 7 | Mucopolysaccharidosis VII (Sly syndrome) | Autosomal recessive |
| *HEXA* | 15 | GM2 gangliosidosis type I (Tay-Sachs disease) | Autosomal recessive |
| *HEXB* | 5 | GM2 gangliosidosis type 2 (Sandhoff disease) | Autosomal recessive |
| *HGSNAT* | 8 | Mucopolysaccharidosis IIIC (Sanfilippo syndrome C) | Autosomal recessive |
| *HYAL1* | 3 | Mucopolysaccharidosis IX | Autosomal recessive |
| *IDS* | X | Mucopolysaccharidosis II (Hunter syndrome) | X-linked recessive |
| *IDUA* | 4 | Mucopolysaccharidosis I (Hurler, Scheie, and Hurler/Scheie syndromes) | Autosomal recessive |
| *LAMP2* | X | Danon disease | X-linked dominant |
| *LIPA* | 10 | Wolman disease<br>Cholesteryl ester storage disease | Autosomal recessive |
| *MAN2B1* | 19 | α-Mannosidosis | Autosomal recessive |
| *MANBA* | 4 | β-Mannosidosis | Autosomal recessive |
| *MCOLN1* | 19 | Mucopolipidosis IV | Autosomal recessive |
| *NAGA* | 22 | Schindler disease types I and II (Kanzaki disease) | Autosomal recessive |
| *NAGLU* | 17 | Mucopolysaccharidosis IIIB (Sanfilippo syndrome B) | Autosomal recessive |
| *NEU1* | 6 | Sialidosis | Autosomal recessive |
| *NPC1* | 18 | Niemann–Pick type C disease | Autosomal recessive |
| *NPC2* | 14 | Niemann–Pick type C disease | Autosomal recessive |
| *PPT1* | 1 | Neuronal ceroid lipofuscinosis 1 (infantile NCL) | Autosomal recessive |
| *PSAP* | 10 | Gaucher disease<br>Metachromatic leukodystrophy | Autosomal recessive |
| *SGSH* | 17 | Mucopolysaccharidosis IIIA (Sanfilippo syndrome A) | Autosomal recessive |
| *SMPD1* | 11 | Niemann–Pick disease type A and B | Autosomal recessive |
| *SUMF1* | 3 | Multiple sulfatase deficiency | Autosomal recessive |
| *TPP1* | 11 | Neuronal ceroid lipofuscinosis 2 (Classic late-infantile NCL) | Autosomal recessive |

\*Inheritance patterns based on the information provided in the Online Mendelian Inheritance in Man database (https://www.omim.org/).
HGNC, HUGO Gene Nomenclature Committee.

We selected PPVs based on three different measures to determine their pathogenicity: (1) predicted mutational effects on the sequence and expression of transcripts and proteins, (2) clinical and experimental evidence obtained from the curated variant databases such as ClinVar, Human Gene Mutation Database (HGMD), and locus-specific mutation databases (LSMDs) and the medical literature, and (3) *in silico* prediction of mutational effects on protein function (Methods). Assuming that variants with a population allele frequency (AF) of ≥0.5% are extremely unlikely to cause LSDs, we excluded variants with an average AF between the Pan-Cancer and 1000 Genomes cohorts higher than this threshold during the PPV selection process. Using an automated algorithm-based approach, a total of 432 PPVs were selected in 41 genes; no PPV was identified in *LAMP2* (Supplementary Fig. 2a and Supplementary Table 2). The selected PPVs were grouped into three tiers with partial overlaps, each tier corresponding to each of the three selection criteria (Fig. 1d).

Overall, PPV prevalence was 20.7% in the Pan-Cancer cohort, which was significantly higher than the 13.5% PPV prevalence of the 1000 Genomes cohort (odds ratio, 1.67; 95% confidence interval, 1.44-1.94; P=8.7×10^−12^; Fig. 2a). This association remained significant after adjustment for population structure (odds ratio, 1.44; 95% confidence interval, 1.22-1.71; P=2.4×10^−5^). The odds ratio for cancer risk was higher in individuals with a greater number of PPVs (P=7.3×10^−12^), and this tendency was broadly consistent when the analysis was restricted to individual tiers, although some tier-specific results did not reach statistical significance (Fig. 2a). For comparison, we examined the prevalence of rare synonymous variants (RSVs) with an average AF between the Pan-Cancer and 1000 Genomes cohorts of <0.5% and found no difference between the two cohorts after adjustment for population structure, indicating that the enrichment of PPVs in the Pan-Cancer cohort was not likely due to batch effects (Fig. 2b). The gene-specific prevalence of PPVs and RSVs in the Pan-Cancer and 1000 Genomes cohorts is shown in Supplementary Figs. 2b and 2c, respectively. The results demonstrated that PPVs were relatively more abundant in the Pan-Cancer cohort versus the 1000 Genomes cohort with respect to the abundance of RSVs, for 33 of 42 genes (78.6%; exact binomial test P<0.001).

**Fig. 2.**
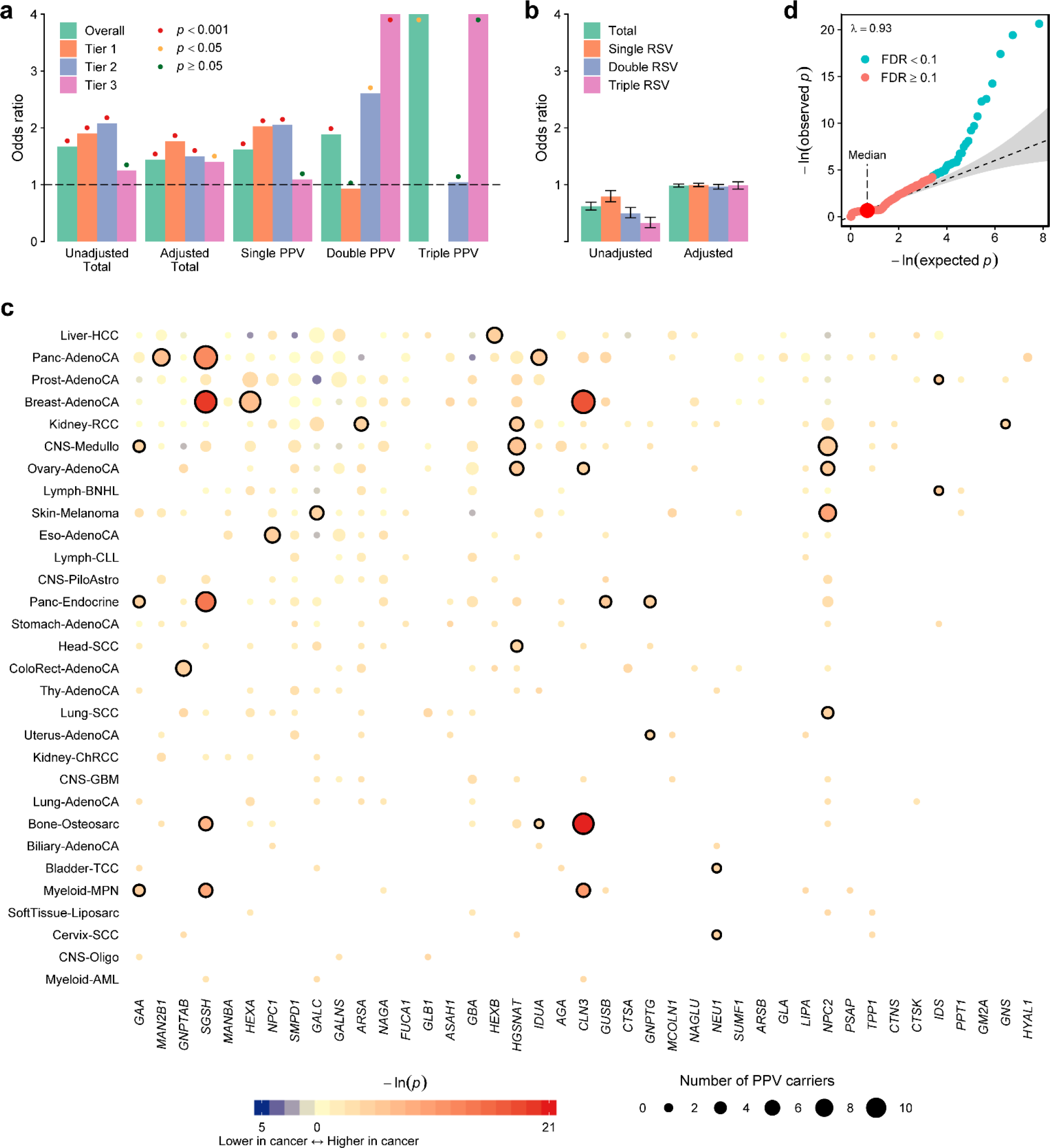
Enrichment of PPVs in cancer patients. **a**, Odds ratios for the prevalence of total PPVs (with or without population adjustment) or PPVs belonging to each of three tiers in the Pan-Cancer versus 1000 Genomes cohorts. Odds ratios for the prevalence of single, double, and triple PPV carriers (individuals carrying one, two, or three PPVs, respectively) are also presented without population adjustment. Odds ratios for double and triple carriers of tier 3 PPVs and triple carriers of total PPVs are 7.54, infinite, and 7.4, respectively, with the corresponding bars cut off at the top edge of the plot. **b**, Odds ratios for the prevalence of RSVs analyzed in the same manner as for PPVs. Error bars indicate 95% confidence intervals. **c**, SKAT-O association between 30 major histological types of cancer (>15 patients per type) and PPVs in each LSD gene. The area of each dot is proportional to the number of PPV carriers for the corresponding cohort-gene pair. Significantly associated cohort-gene pairs at the 0.1 FDR threshold are encircled by bold rings. Cohorts are shown in descending order according to the number of patients they include (top to bottom), and genes are shown in descending order according to the number of unique PPVs they contain (left to right). Abbreviations of histological diagnoses are defined in Supplementary Table 1. **d**, Quantile-quantile plot of P-values derived from SKAT-O analyses. A group-based inflation factor (λ) is displayed at the top left-hand corner (Methods). Gray shading indicates the 95% confidence interval. Each dot in this plot corresponds to each dot shown in **c**.

### Association of PPVs with specific cancer types

Among the 30 major histological types of cancer (>15 individuals per cancer type), the PPV prevalence ranged from 8.8% to 48.6%, with significantly higher values in seven histological types of cancer than in the 1000 Genomes cohort (Supplementary Fig. 3a). Results of tier-based analyses were broadly consistent (Supplementary Figs. S3b–d). In contrast, RSV prevalence showed much less variation across cohorts and was higher in the 1000 Genomes cohort than in any cancer cohort (Supplementary Fig. 3e), reflecting the more heterogeneous nature of ancestry (Fig. 1b) and the resulting higher genetic polymorphism in the 1000 Genomes cohort. Analysis using the optimal sequence kernel association test (SKAT-O) method, adjusted for population structure (Methods), unveiled 37 significantly associated cancer-gene pairs and four genes (*GBA*, *SGSH*, *HEXA*, and *CLN3*) with a pan-cancer association (Fig. 2c and Supplementary Fig. 2b and Supplementary Table 3). Overall, 19 cancer types were significantly enriched for PPVs in at least one LSD gene, and PPVs in 18 genes were associated with at least one cancer type. We observed no evidence of systematic inflation of test statistics (Fig. 2d).

### PPV prevalence in the Pan-Cancer and ExAC cohorts

We sought to validate the findings of the SKAT-O analysis using the ExAC cohort as an independent control. For this purpose, we focused on (1) eight cancer cohorts that showed significantly higher PPV prevalence than the 1000 Genomes cohort (Supplementary Fig. 3a) and (2) ten PPV groups that were significantly enriched in the Pan-Cancer cohort or three or more histological cancer subgroups compared to the 1000 Genomes cohort (Fig. 2c and Supplementary Fig. 2b). As shown in Supplementary Fig. 4a, PPV prevalence was higher in all tested cancer cohorts than in the ExAC cohort, and the association was significant for the Pan-Cancer, pancreatic adenocarcinoma, medulloblastoma, pancreatic neuroendocrine carcinoma, and osteosarcoma cohorts. In addition, all tested PPV groups except *GBA* were more prevalent in the Pan-Cancer cohort than in the ExAC cohort, and six were significantly enriched in cancer patients (Supplementary Fig. 4b).

### Variant-specific enrichment of PPVs in cancer patients

Although our cohorts were underpowered for detection of variant-specific cancer association for such rare variants as PPVs, some results deserve attention. Among all 432 PPVs identified in the Pan-Cancer and 1000 Genomes cohorts (Fig. 1d), a splicing variant in *NPC2*, rs140130028 (ENST00000434013:c.441+1G>A), was most strongly associated with various histological types of cancer including medulloblastoma (P=0.008), ovarian adenocarcinoma (P=0.022), cutaneous melanoma (P=0.003), and lung squamous cell carcinoma (P=0.019; Supplementary Fig. 5). Inactivating mutations of *NPC2* cause Niemann-Pick type C disease, which typically presents as progressive neurological abnormalities. The relationship between the Niemann-Pick type C disease and medulloblastoma was implied by a structural homology of NPC1 with Patched transmembrane protein, a tumor suppressor that is regulated by Hedgehog signaling and involved in the development of medulloblastoma when inactivated by loss-of-function mutations.^16, 17^ Vismodegib, a downstream Hedgehog signaling inhibitor, showed promising antitumor activity in animal models, leading to evaluation of this agent in clinical trials for the treatment of medulloblastoma.^18, 19^ Nonetheless, no study to date has provided direct evidence linking medulloblastoma to mutations causing Niemann-Pick type C disease. Results of our study, therefore, provide the first genetic evidence of the tumorigenic potential of inactivating *NPC2* mutations.

Another example, rs145834006—a 3′ UTR variant in *IDS* that was significantly associated with downregulated gene transcription (P=1×10^−5^; false discovery rate [FDR] = 0.07; Supplementary Fig. 6)—showed a strong association with non-Hodgkin B-cell lymphoma (P=2.2×10^−4^). This finding was in accordance with the significant SKAT-O association between *IDS* PPVs and non-Hodgkin B-cell lymphoma (P=0.005; FDR=0.068; Fig. 2c). The relatively high *IDS* expression in lymphoid tissue implies an essential role of the protein encoded by this gene in lymphoid organ function (Supplementary Fig. 7). Collectively, our results generate a plausible hypothesis of the lymphomagenic property of *IDS* loss-of-function mutations that warrants confirmation in larger lymphoma cohorts and functional studies.

### Age at diagnosis of cancer according to PPV carrier status

The age at diagnosis of cancer across 28 major clinical cancer cohorts (corresponding to 30 major histological types that included 15 or more patients; information on age at diagnosis was not available for patients with osteosarcoma; patients with pilocytic astrocytoma and oligodendroglioma were combined into a single clinical cohort; see Methods) is shown in Fig. 3a. To examine whether cancer occurred earlier in PPV carriers than in wild-type individuals, we first compared the age at diagnosis according to PPV carrier status in the Pan-Cancer cohort and in six clinical cancer subgroups that showed significant SKAT-O association with PPVs (Fig. 3b). The median age at diagnosis of cancer was numerically lower in PPV carriers in all evaluated cohorts, and the difference was significant in the following cohorts: Pan-Cancer (median age, 59 versus 61 years; P=0.002), pancreatic adenocarcinoma (median age, 61 versus 68.5 years; P<0.001), and chronic myeloid disorder (median age, 45.5 versus 58.5 years; P=0.044). We next compared the age at diagnosis of cancer between carriers and non-carriers of PPVs that belonged to each PPV group that was significantly enriched in the Pan-Cancer cohort or three or more cancer types compared to the 1000 Genomes cohort—same criteria were used for the validation of SKAT-O results with the ExAC cohort as an independent control, as described above—among the Pan-Cancer cohort. As shown in Fig. 3c, carriers of PPVs that belonged to tier 1, tier 3, *HGSNAT*, *CLN3*, and *NPC2* had a significantly earlier onset of cancer compared to wild-type individuals. Moreover, the PPV load (number of PPVs per individual) showed a consistent negative linear correlation with age at diagnosis of cancer across all histological types and PPV groups evaluated, and the correlation was significant in the Pan-Cancer and pancreatic adenocarcinoma cohorts (Figs. 3d and 3e). Exploratory analysis across all cancer types and genes revealed earlier cancer onset in PPV carriers for five additional cancer-gene pairs (Fig. 3f), three of which (pancreatic adenocarcinoma-*MAN2B1*, cutaneous melanoma-*NPC2*, and chronic myeloid disorder-*SGSH*) were in concordance with the SKAT-O results (Fig. 2c).

**Fig. 3.**
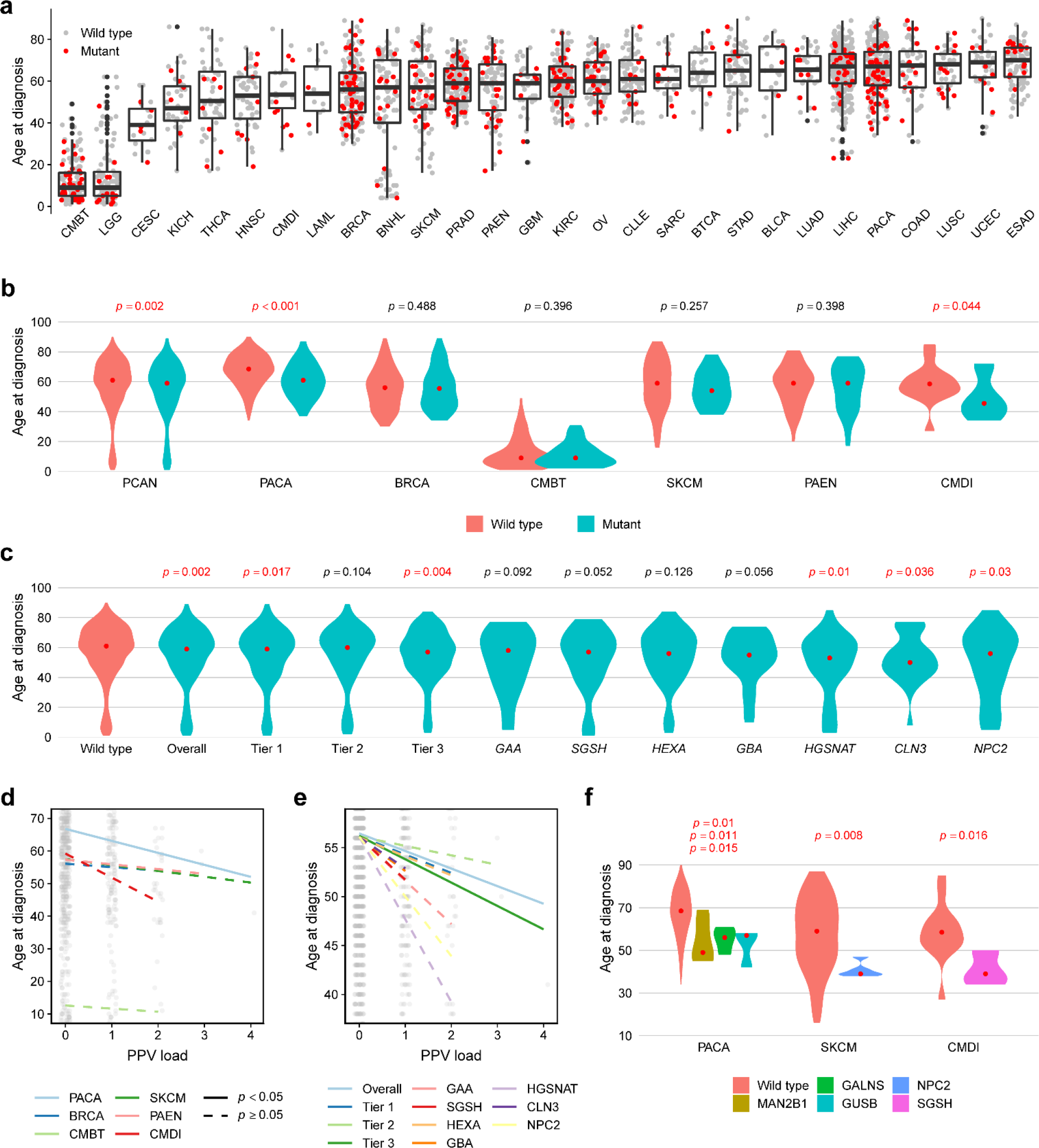
Age at diagnosis of cancer. **a**, Age at diagnosis of cancer across 28 major clinical cancer cohorts. Patients are represented by red (PPV carrier) or gray (non-carrier) dots. Boxes encompass the 25th through 75th percentiles, the horizontal bar represents the median, and the upper and lower whiskers extend from the upper and lower hinges to the largest and smallest values no further than 1.5 × interquartile range from the hinges, respectively. Data beyond the end of whiskers are plotted individually. **b**, Age at diagnosis of cancer in carriers and non-carriers of PPVs in the Pan-Cancer cohort and six clinical cancer subgroups that showed significant SKAT-O association with PPVs. **c**, Age at diagnosis of cancer according to the carrier status of 11 PPV groups significantly associated with the Pan-Cancer cohort or more than two histological cancer subgroups in the SKAT-O analysis. **d**,**e**, Linear correlations between the PPV load and age at diagnosis of cancer in six clinical cancer subgroups shown in **b**(**d**) and in the Pan-Cancer cohort for each of 11 PPV groups shown in **c**(**e**). In **d** and **e**, each dot represents a single patient. Simple linear regression was performed for each cohort in **d**, and linear regression adjusted for cancer histology was performed for each group of PPVs in **e**to draw the regression line and test for statistical significance. As plots in **d** and **e**are magnified to clearly distinguish between regression lines, not all patient dots are included within the plotted area. **f**, All cancer-gene pairs in which age at diagnosis of cancer differs significantly according to the PPV carrier status. In **b**, **c**, and **f**, P-values derived from one-sided Wilcoxon rank sum tests are shown above each violin plot. The vertically aligned P-values from top to bottom for PACA in **f** correspond to the three genes displayed from left to right, respectively. The red dot in each violin plot represents the median. See Supplementary Table 1 for abbreviations of clinical cancer cohorts.

### Differential somatic mutation and gene expression patterns of pancreatic adenocarcinoma from PPV carriers

We sought to determine whether differentiating patterns of somatic mutations and gene expression underlie the oncogenic processes triggered by PPVs in pancreatic adenocarcinoma, for which both the SKAT-O analysis and comparison of age at diagnosis of cancer according to PPV carrier status produced consistent results (Figs. 2c, 3b, 3d, and 3f and Supplementary Figs. 3a–d and 4). We first compared the somatic mutational landscape between tumors from PPV carriers (n=55) and non-carriers (n=177). The 50 most frequently mutated genes in each group are shown in Supplementary Fig. 8. The five top-ranked genes were common in both groups (*KRAS*, *TP53*, *SMAD4*, *CDKN2A*, and *TTN*), and the first four of these were in agreement with previous genome sequencing studies of pancreatic adenocarcinoma.^20,21^ Non-silent mutation burden was similar between groups (mean 57.1 versus 56.3 mutations per tumor for PPV-associated versus PPV-unrelated cases, respectively; P=0.9). Mutational signature also did not differ according to the PPV carrier status (P≥0.05 for all signatures; Supplementary Fig. 9).

Differentially expressed gene (DEG) analysis of pancreatic adenocarcinoma samples with available RNA-Seq data (n=55; 8 from carriers and 47 from non-carriers of PPVs) revealed 287 gene upregulations and 221 downregulations in tumors from PPV carriers compared to those from wild-type individuals (Figs. 4a–d and Supplementary Table 4). Pathway-based analysis with the generally applicable gene set enrichment (GAGE) method identified 63 pathways significantly altered by PPV carrier status (Fig. 4e and Supplementary Fig. 10). Remarkably, these pathways included at least six among 13 core signaling pathways that have been shown to be recurrently perturbed in pancreatic cancer: Ras signaling, Wnt signaling, axon guidance, cell cycle regulation, focal adhesion, cell adhesion, and ECM-receptor interaction pathways.^21, 22^ In addition, our data suggested that deleterious mutations in LSD genes can provoke perturbations in neurodegenerative disease pathways involved in the development of Parkinson disease, Alzheimer disease, and Huntington disease, all of which have been reported to occur frequently in LSD patients.^1^ The glycerophospholipid metabolism pathway was also identified, indicating that altered gene expression and nonsense-mediated decay might have contributed to lysosomal dysfunction in PPV carriers.

**Fig. 4.**
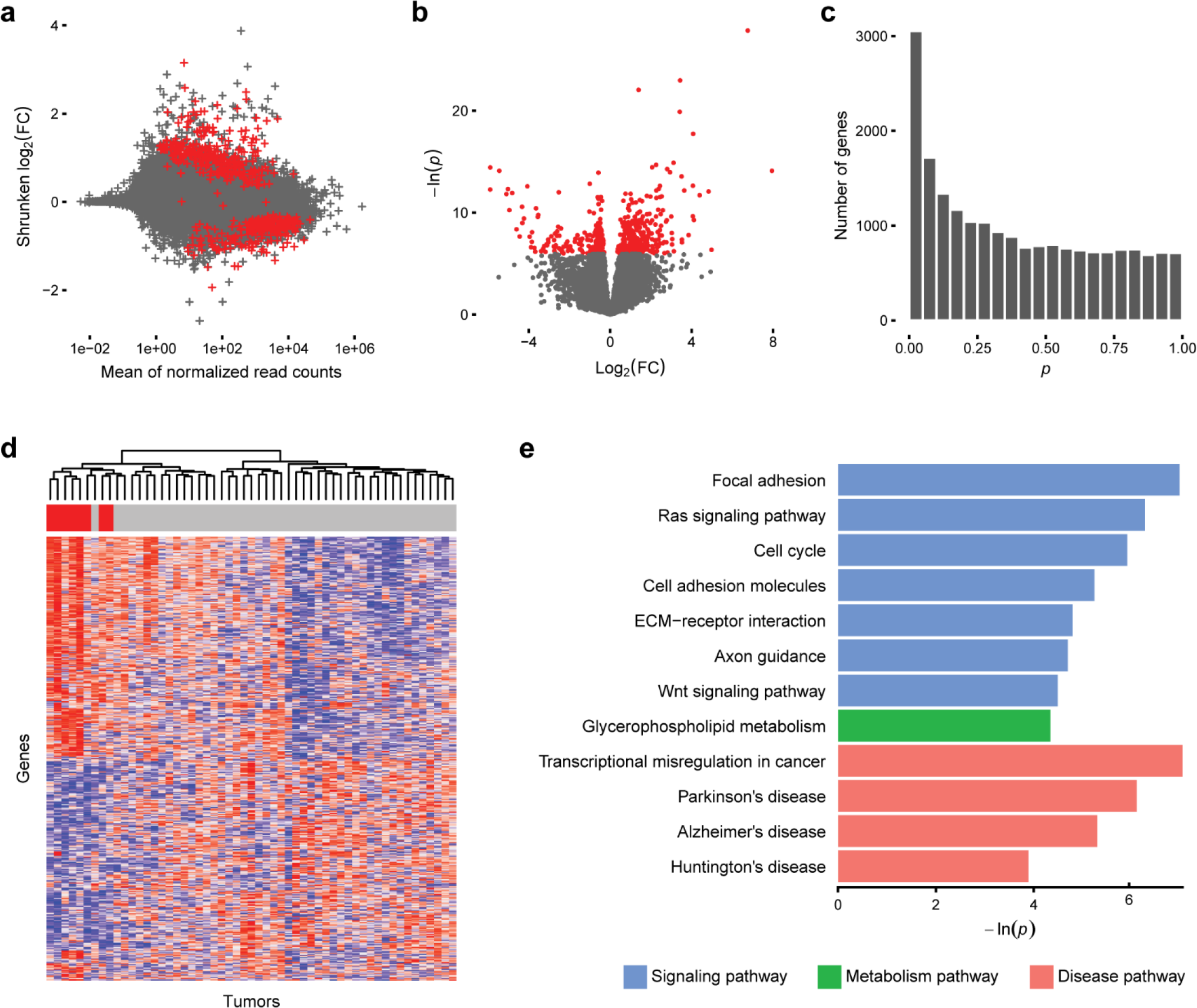
Differentially expressed genes and pathways in pancreatic adenocarcinoma from PPV carriers versus non-carriers. **a-c**, DEG analysis reveals 287 gene upregulations and 221 downregulations in PPV-associated pancreatic adenocarcinoma. In **a** and **b**, genes with FDR<0.1 are shown in red. FC, fold change. In **c**, the histogram of P-values shows a peak frequency below 0.05, demonstrating the existence of up-or downregulated genes. **d**, Heatmap showing the relative expression of genes significantly up-or downregulated at the 0.1 FDR threshold in tumors from PPV carriers versus non-carriers, labeled with red and gray bars under the dendrogram, respectively. We ranked the samples according to the FPKM-UQ-normalized read counts for each gene and used the rank numbers for color mapping in order to standardize the visual contrast across genes. Samples are ordered as columns by hierarchical clustering based on the Euclidean distance and complete linkage. Genes are ordered as rows in the same manner (dendrogram not shown). High and low relative expression is indicated by progressively more saturated red and blue colors, respectively. **e**, KEGG pathways that are significantly altered in tumors from PPV carriers compared to those from non-carriers. Only pathways of particular interest discussed in the text are shown. All pathways with FDR <0.1 are shown in Supplementary Fig. 14. ECM, extracellular matrix.

### Supportive data

Additional data including the pathogenic variant detection capability of the tier 3 PPV selection criterion, PPV-to-synonymous variant prevalence ratios, and population-specific prevalence of PPVs in *SGSH* are provided in Supplementary Information.

## Discussion

In the present study, we showed that potentially pathogenic germline mutations in LSD genes, identified based on three different pathogenicity criteria, were significantly enriched in cancer patients across a wide range of histological types. Our aggregate rare-variant association analysis approach enabled detection of rare variant enrichment both in the Pan-Cancer cohort and in a histology-dependent manner, which would have been undetectable by using conventional variant-wise association analysis methods. Analysis by subgrouping of PPVs into three tiers based on different selection criteria, validation with an independent control cohort, and comparison of the results with those obtained from synonymous variants with matched AF showed broadly consistent results, corroborating the findings of our study. The genetic association was further supported by the significant difference in age at diagnosis of cancer observed in carriers versus non-carriers of PPVs in the Pan-Cancer cohort as well as at least two clinical subgroups of cancer patients.

The lysosome is involved in a variety of cellular functions other than biomolecule catabolism, such as intracellular signaling, nutrient sensing, cellular growth regulation, plasma membrane repair, and phagocytosis.^15^ The diverse roles of lysosomes underlie the complex and heterogeneous phenotypes of LSDs which can involve almost any organ.^1^ It has long been evident that patients with Gaucher disease are at markedly increased risk of malignancy, especially multiple myeloma with the risk estimated at approximately 50-fold.^23^ However, despite the largely shared pathogenesis, the relationship between most other LSDs and cancer has been largely unexplored because of the rarity and phenotypic heterogeneity of each LSD. The wide spectrum of tumor histologies and LSD genes covered in our study enabled elucidation of numerous cancer-gene pairwise associations most of which had previously been unknown.

From the SKAT-O analyses, we identified four genes that showed a significant pan-cancer association; among those, *SGSH* and *CLN3* were strongly associated with five and four cancer types, respectively. *SGSH* encodes sulfamidase, a lysosomal hydrolase that degrades heparan sulfate. Deficiency of sulfamidase leads to Sanfilippo syndrome A (mucopolysaccharidosis IIIA), which is characterized by progressive mental and behavioral deterioration that typically presents in childhood. However, an adult-onset disease that presents primarily with visceral manifestations without neurological abnormality has also been reported.^24^ A recent *in vivo* study suggested a crucial role of oxidative stress in the pathobiology of Sanfilippo syndrome A.^25^ Since the oxidative stress is a key mediator of cancer cell growth, invasiveness, and angiogenesis,^4^ inherited *SGSH* mutations may contribute to an elevated cancer risk via persistent cellular exposure to oxidative stress, a plausible hypothesis that should be confirmed in future functional studies.

CLN3 is a late endosomal and lysosomal transmembrane protein, and its defect causes classic juvenile neuronal ceroid lipofuscinosis (CLN3 disease). In CLN3 disease, impaired trafficking of galactosylceramide to the plasma membrane promotes the generation of proapoptotic ceramide and subsequent activation of caspases, which in turn accelerates apoptosis.^14^ In line with its control over apoptosis, CLN3 also regulates cancer cell growth, and its therapeutic implication has been suggested.^26^ Therefore, results of our study warrant future investigation of this protein as a therapeutic target for the treatment of various types of cancer.

Almost 5 to 10 percent of pancreatic cancer patients are diagnosed before the age of 50.^27^ For these patients, positive family history is a strong risk factor, indicating the presence of inherited risk variants.^28^ Indeed, many pancreatic cancer-predisposing mutations have been identified in genes involved in the genome maintenance and double-strand DNA break repair (e.g., *BRCA1/2* and *PALB2*). However, in a majority of the early-onset pancreatic cancer patients, the genetic cause remains unclear.^29^ In our histology-specific analysis, patients with pancreatic adenocarcinoma showed a strong association with PPVs in several LSD genes and had a significantly earlier onset of cancer, motivating us to evaluate differential patterns of somatic mutations and gene expression in this histological subset. The DEG analysis revealed many genes up-or downregulated in PPV carriers, and GAGE analysis provided novel insights into the biological processes that might be involved in pancreatic carcinogenesis in these patients. Remarkably, many of the altered pathways identified in the GAGE analysis were previously implicated in pancreatic cancer development in transcriptome and exome sequencing studies.^21, 22^ The somatic mutation burden and signatures, in contrast, were comparable between the carriers and non-carriers of PPVs. Overall, the results of our study suggest that transcriptional misregulation is a key mediator of pancreatic carcinogenesis triggered by PPVs.

Decades of research on tumor suppressor genes involved in hereditary cancer predisposition syndromes have proposed a continuum model of tumor suppression, emphasizing the crucial impact of a subtle change in expression of tumor suppressor genes.^30^ Given the rarity of individual PPVs, almost all PPV carriers observed in our study were heterozygous. Therefore, the dosage effect model may be useful in explaining the mechanisms involved in oncogenic contributions of LSD gene mutations, which has already been implied by results of a previous study.^9^

Several limitations of this study require careful acknowledgment. As we did not process the raw sequence data but used variant call sets produced by independent research consortiums, the possibility of batch effects cannot be excluded, even considering the similarity in pipelines used to generate each dataset.^10–12^ Second, although the ExAC cohort served as a large-scale validation control set, we could not adjust for the population structure in association analysis using this cohort because the individual-level genotype data were not accessible. The ExAC cohort has a similar population composition to the Pan-Cancer cohort, with almost 70% of the entire cohort comprised of Americans and Europeans,^12^ but this similarity does not remove the need for population correction. Third, an independent cancer cohort that is sufficiently powered for analyzing such rare variants as PPVs was not available for external validation. The PCAWG project currently represents the largest cancer genome analysis effort harmonizing data from different sources into a single, contemporary state-of-the-art pipeline. Therefore, validation of our findings in an external cancer patient cohort will be possible only with additional cancer patient genomes sequenced and harmonized with the existing database in the future. Finally, hematological malignancies such as myeloma, the most widely known LSD-associated cancer, were poorly represented in the Pan-Cancer cohort, and the numbers of patients with individual cancer types were not sufficiently large to draw reliable histology-specific conclusions.

From a therapeutic perspective, LSD genes are attractive targets because of the mechanistically intuitive nature of the enzyme replacement and substrate reduction therapies. The enzyme replacement therapy has already been approved for at least seven types of LSD.^31^ Other promising approaches include pharmacological chaperones, gene therapy, and compounds that ‘read through’ the early stop codon introduced by nonsense mutations.^31^ Although it is unclear whether preemptive treatment can prevent or delay long-term complications of LSDs such as cancer, our findings make it promising to harness these sophisticated LSD therapies for preventing cancer in carriers of inactivating germline mutations in LSD genes.

In conclusion, the present study provides a comprehensive landscape of association between potentially pathogenic germline mutations in LSD genes and cancer. Investigating the crosstalk between treatable metabolic diseases and cancer is crucial since it can build the basis for precision cancer prevention. Diverse and increasingly sophisticated therapeutic options to restore lysosomal functions are currently available or being developed. Future clinical trials of these agents guided by individuals’ mutation profiles may pave a new path toward personalized cancer prevention and treatment.

## Methods

### Data sources

We downloaded germline and somatic (tumor) variant datasets for SNVs and indels of the Pan-Cancer cohort as variant call format (VCF) and mutation annotation format (MAF) files, respectively, from the sftp server of the PCAWG project (http://dccsftp.nci.nih.gov/pancan/). The germline variant call sets encompassed all 2,834 PCAWG donors and were produced using the DKFZ/EMBL pipeline. The tumor somatic MAF file contained data of 2,583 whitelist samples (only one representative tumor from each multi-tumor donor) and were generated by the PCAWG consensus strategy consolidating outputs from the Sanger, Broad, DKFZ/EMBL, and MuSE pipelines for SNVs and from the SMuFin, DKFZ, Sanger, and Snowman pipelines for indels. Pass-only variants were used for the analysis. We downloaded tumor RNA-Seq data as both raw and normalized read count matrices of protein-coding genes via Synapse (https://www.synapse.org/#!Synapse:syn3104297). Read alignment was carried out using TopHat2, counted using the htseq-count script from the HTSeq framework version 0.6.1p1 against the reference General Transfer Format of GENCODE release 19, and normalized using the FPKM-UQ normalization technique.^32^ We downloaded the clinical and histological annotation sheets from the PCAWG wiki page (https://wiki.oicr.on.ca/pages/) both in version 9 (generated on November 22, 2016 and August 21, 2017, respectively).

As the primary control cohort, we downloaded the individual-level genotype data of SNVs and indels for 2,504 individuals from the 1000 Genomes project phase 3 (1000 Genomes cohort) as VCF files (http://ftp.1000genomes.ebi.ac.uk/vol1/ftp/phase3).^11^ In addition, population-level AF data of SNVs and indels for 53,105 unrelated individuals from the ExAC release 1.0 (ExAC cohort), excluding TCGA subset, were downloaded for use as an independent validation control (http://ftp.broadinstitute.org/pub/ExAC_release/release1).^12^

### Quality assessment and control

Quality assessment of all PCAWG sequence data was carried out according to three-level criteria (library, sample, and donor levels) to determine whether to include each donor and RNA-Seq aliquot in the study or not. This multi-level quality control process was necessary since individual donors could have multiple samples, and individual samples could have multiple libraries. As a rule, a sample was blacklisted if all of its libraries were of low quality, and whitelisted if all of its libraries were of high quality. Similarly, a donor was blacklisted if all associated samples were blacklisted, and whitelisted if all associated samples were whitelisted. Samples and donors that were neither blacklisted nor whitelisted were graylisted. Only whitelisted individuals and samples were included in the present study (2,583 tumor-normal pair genomes and 1,094 RNA-Seq samples). Quality control criteria for each level of assessment are detailed in the PCAWG marker paper.^10^

### Consolidation of the Pan-Cancer cohort

The original PCAWG project covered 2,834 individuals encompassing 40 major cancer types as part of the ICGC, which included 76 projects and 21 primary organ sites.^10^ Among those, we prioritized 2,583 whitelisted patients who satisfied the multi-level quality control criteria described above. Sixteen patients with a histological diagnosis indicating a benign bone neoplasm, such as chondroblastoma, chondromyxoid fibroma, osteofibrous dysplasia, and osteoblastoma, were excluded, leaving 2,567 patients in the final Pan-Cancer cohort. Nine patients who had multiple tumor specimens were associated with more than one histological diagnosis: eight patients with both myeloproliferative neoplasm and acute myeloid leukemia and one patient with both hepatocellular carcinoma and cholangiocarcinoma. For consistency in the histology-specific analysis, the first eight patients were classified as acute myeloid leukemia and the ninth patient as cholangiocarcinoma. To analyze the age at diagnosis of cancer, we combined multiple histological cohorts that shared similar clinicopathologic characteristics into a single clinical cohort (e.g., breast invasive ductal, lobular, and micropapillary carcinomas were classified as breast cancer [BRCA], and myeloproliferative neoplasm and myelodysplastic syndrome as chronic myeloid disorder [CMDI]; see Supplementary Table 1). Among the 2,567 patients, only 1,075 had whitelisted tumor RNA-Seq data. Since 19 patients contributed more than one tumor specimen, RNA-Seq data were available for 1,094 tumors.

### Gene selection and variant interpretation

Of the genes involved in lysosomal functions that include substrate hydrolysis, post-translational modification of hydrolases, intracellular trafficking, and enzymatic activation, we selected 42 genes that were previously implicated in the development of LSD via comprehensive literature review.^1,8,13–15^ The genomic loci of the selected genes based on the GRCh37/hg19 human reference genome assembly were screened for all germline SNVs and indels in each normal VCF file. Variants were identified based on the GENCODE release 19 gene model (https://www.gencodegenes.org/releases/19.html). We carried out functional annotation using both ANNOVAR and Variant Effect Predictor version 85 and cross-checked and manually curated the outputs to achieve the most appropriate characterization of each identified variant.^33, 34^ From this point, our analysis focused on variants within protein-coding regions, splice donor and acceptor sites within two base pairs to the intron side from the exon-intron junctions (GT-AG conserved sequence), and 5′ and 3′ UTRs. Variants were classified into ten non-overlapping categories according to the predicted consequence type on transcripts or proteins: missense, start-loss, stop-gain, stop-loss, synonymous, frameshift indel, non-frameshift indel, splicing, and 5′ and 3′ UTR variants. When a variant was associated with more than one consequence type depending on transcript isoforms, it was classified into the most functionally disruptive category (e.g., protein-truncating rather than missense, and missense rather than UTR or synonymous). For example, rs373496399 (NC_000017.10:g.78184457G>A) could be either a missense or 3′ UTR variant depending on the transcript isoform and was classified as missense. By this way, each variant belonged to a unique functional class that was used for subsequent analysis. *In silico* prediction of the mutational effect on protein function was carried out by using 19 distinct computational algorithms with the use of dbNSFP version 3.3 (Supplementary Fig. 11).^35—52^

### PPV selection

The prevalence of individual LSDs ranges from one per tens of thousands to one in millions of live births, and considerable allelic heterogeneity exists.^53–55^ Therefore, a single variant with a population AF ≥0.5% is extremely unlikely to be causative, even considering the possibility of underdiagnosis. A recent analysis of the prevalence of known Mendelian disease variants using >60,000 exomes sequenced suggested that a substantial proportion of variants with AF >1% were, in fact, benign or functionally neutral, highlighting the importance of filtering PPVs based on their frequency in a sufficiently large reference population.12 On this theoretical basis and our data showing that deleterious variants were rare, mostly with an AF of <0.5% (Supplementary Fig. 12), we excluded variants with an average AF between the Pan-Cancer and 1000 Genomes cohorts of ≥0.5% during the PPV selection process.

We examined the curated databases ClinVar, HGMD, and LSMDs and extensively reviewed the medical literature to identify LSD-causing mutations (Supplementary Table 5). We initially classified variants into five non-overlapping categories, as proposed by the American College of Medical Genetics and Genomics (ACMG) and Association for Molecular Pathology (AMP) based on the curated clinical significance information in ClinVar.^56^ In case of variants that belonged to more than one pathogenicity category, priority was assigned to the category associated with stronger evidence, hence ‘benign’ rather than ‘likely benign,’ and ‘pathogenic’ rather than ‘likely pathogenic.’ When interpretations indicating both pathogenic (‘pathogenic’ or ‘likely pathogenic’) and benign (‘benign’ or ‘likely benign’) directions of effect coexisted for a single variant, or no pathogenicity interpretation was provided in standard terminology, data in HGMD and LSMDs along with supporting evidence obtained from direct literature survey were reviewed to determine the most relevant functional category of the variant according to the ACMG and AMP guideline.

As the role of microRNA in carcinogenesis has been spotlighted in recent years,^57, 58^ researchers have identified many SNVs in 3′ UTR microRNA-binding sites that were involved in the increased or decreased cancer risk via altered expression of gene products.^59–63^ Although much less identified, 5′ UTRs also contain binding motifs for microRNAs, and their sequence variation affects messenger RNA (mRNA) stability.^64, 65^ Since UTR variants can create or destroy a microRNA-binding motif that regulates gene expression and mRNA degradation, the biological consequence of UTR variants can be reflected in the change in transcript abundance in relevant tissues.^66, 67^ Therefore, we analyzed RNA-Seq read count data to identify UTR variants associated with significantly decreased expression of the corresponding genes. Among the 3,192 unique UTR variants with mean AF <0.5% between the Pan-Cancer and 1000 Genomes cohorts, 795 and 2,397 were present in 5′ and 3′ UTRs, respectively. We compared the tissue mRNA abundance after variance-stabilizing transformation of read counts between UTR variant carriers and non-carriers for each gene, using linear regression.^68^ Because the expression level of each LSD gene varied considerably across cancer types (e.g., *IDS* shown in Supplementary Fig. 7), the regression model was adjusted for cancer histology. As a result, only one 3′ UTR variant in *IDS*, rs145834006 (ENST00000340855:c.*3950A>G), reached statistical significance at the 0.1 FDR threshold (Supplementary Fig. 6).

After inspecting all information obtained from the above processes, we selected PPVs that were highly likely to cause LSD by using three positive selection criteria (Fig. 1d). Tier 1 included all frameshift indels, start-loss variants, stop-gain variants, splicing variants, and a UTR variant associated with significant downregulation of the corresponding gene (rs145834006). Thus, most of these variants were loss-of-function in principle. Tier 2 included variants classified as ‘pathogenic’ or ‘likely pathogenic’ based on the information obtained from ClinVar and relevant medical literature, disease-causing mutations in HGMD (designated as ‘DM’ in the database), and pathogenic mutations ascertained via LSMDs. Of the variants without curated pathogenicity information in both ClinVar and HGMD (i.e., with unknown clinical significance), those predicted to be functionally deleterious by all of the 19 separate *in silico* prediction tools were classified into tier 3. The score threshold of each tool for classifying a variant as deleterious or benign was set at the provided default when available, or the median of all evaluated variants otherwise. Because some variants (especially those in the noncoding regions and indels) were not successfully annotated by all of the 19 tools, only available scores were used in such cases.

### PPV-cancer association analysis using the Pan-Cancer and 1000 Genomes cohorts

Because our cohorts were underpowered to detect variant-specific associations for such rare variants as PPVs, we performed tier-and gene-based aggregate association analysis using the SKAT-O method with an optimal ρ parameter chosen from a grid of eight points (0, 0.1^2^, 0.2^2^, 0.3^2^, 0.4^2^, 0.5^2^, 0.5, 1), which could be interpreted as a pairwise correlation among the genetic effect coefficients.^69^ The SKAT-O method is robust against the co-existence of pathogenic and benign variants and is thus suitable when no uniform assumption can be made for the genetic effects of variants as in the present study. To examine if the difference in variant calling pipelines used in the PCAWG project and the 1000 Genomes project (batch effects) affected our results, we compared the PPV-to-synonymous variant prevalence ratios between cancer cohorts and the 1000 Genomes cohort using weighted logistic regression. For an exploratory purpose, we also assessed the variant-specific association of PPVs with each type of cancer using logistic regression assuming a multiplicative risk model. All association analyses were adjusted for population structure using the method described below.

### Population structure adjustment

For adjustment of population structure, we carried out a principal component analysis using the individual-level genotype data of tag single nucleotide polymorphisms (tag-SNPs) of the Pan-Cancer and 1000 Genomes cohorts. We first downloaded a list of 1,555,886 candidate tag-SNPs from the phase 3 HapMap ftp server (http://ftp.ncbi.nlm.nih.gov/hapmap/phase_3/). We converted the genomic coordinates of these SNPs into the GRCh37/hg19 framework using the Batch Coordinate Conversion (liftOver) tool (https://genome.ucsc.edu/cgi-bin/hgLiftOver). VCF files from both Pan-Cancer and 1000 Genomes cohorts were merged using the Genome Analysis Toolkit to calculate broad AFs.^70^ We used the VCFtools version 1.13 to extract candidate tag-SNPs with AF ≥5% and ≤50% from the merged VCF, leaving 16,304 SNPs in the aggregate genotype matrix.^71^ Among those, we prioritized the population-stratifying tag-SNPs using the PLINK pruning method.^72^ During this process, we used a recursive sliding-window procedure to exclude SNPs with a variance inflation factor >5 within a sliding window of 50 SNPs, shifting the window forward by 5 SNPs at each step. As a result, we reduced the linkage disequilibrium panels containing multiple correlated SNPs to 10,494 representative tag-SNPs, which were used in the subsequent principal component analysis.

A total of 5,071 principal components (PCs) were obtained by performing principal component analysis against the combined genotype data for the 10,494 tag-SNPs of the Pan-Cancer and 1000 Genomes cohorts. We calculated the correlations of each PC with the binary phenotype (cancer versus normal) and PPV load. Predictably, PC1 and PC2 collectively accounted for more than 11% of the total variance and only these two were significantly correlated with both the binary phenotype and PPV load at the 0.1 FDR threshold (Supplementary Fig. 13a). The remaining 5,069 PCs each accounted for less than 1% of the variance and were correlated with either the phenotype or the PPV load or neither, suggesting that only the two top-ranked PCs were potential confounders of the association between PPVs and cancer (Supplementary Figs. 13b–g). Therefore, we included PC1 and PC2 as covariates in the subsequent association analyses. To examine the possibility of systematic inflation of test statistics, we calculated a group-based inflation factor (*λ*) from the histology-specific SKAT-O results using a previously described method (Fig. 2d).^73^

### RNA-Seq data analysis

We filtered out genes with zero read counts across all tumors from the read count matrices to improve the computational speed. Since the data were generated on the framework of Ensembl gene classification, we converted the Ensembl gene ID to Entrez gene ID using Pathview.^74^ When multiple Ensembl IDs matched to a single Entrez ID, those with the largest variance across all samples were selected while the others were removed from the count matrix. We investigated the differential gene expression patterns between tumors from PPV carriers and non-carriers using DESeq2, after applying the shrinkage estimation of log fold changes and dispersions to improve the stability of the estimates (Fig. 4a).^75^ Before estimating FDRs for DEG results, we performed independent filtering of low-count genes using Genefilter to improve statistical power.^76^

Before the GAGE analysis, we performed variance-stabilizing transformation of raw read counts to achieve homoscedasticity of the count matrix and decrease the influence of genes with an excessively large variation in expression level across samples. The GAGE analysis was based on group-on-group comparisons, which could be controlled by the ‘compare’ argument supported by the ‘gage’ function of the Bioconductor package ‘gage.’ We simultaneously tested for the upregulation and downregulation of gene components constituting each Kyoto Encyclopedia of Genes and Genomes (KEGG) pathway in tumors from PPV carriers compared to those from non-carriers.

### Validation analysis using the ExAC cohort as an independent control

Because the ExAC dataset covered only exonic regions consisting of the GENCODE release 19 coding regions and their flanking 50 base pairs,^12^ we restricted our analysis to coding regions covered in more than half of the ExAC samples (median coverage depth ≥1) in the validation analysis. Coverage depth for the ExAC sequence data was downloaded from the ftp site (ftp.broadinstitute.org/pub/ExAC_release/release1/coverage). We then selected PPVs from the aggregate variant call set of the Pan-Cancer and ExAC cohorts using the same criteria used in the primary analysis of the Pan-Cancer and 1000 Genomes cohorts (Fig. 1d). As a result, we identified 1,267 PPVs: 942 in tier 1 and 475 in tier 2 with 150 overlaps between the two tiers. No tier 3 PPV was identified because the pathogenicity score thresholds used for classifying each variant as deleterious or neutral were set at stricter values than in the primary analysis for some of the 19 *in silico* prediction tools. The changes in thresholds were owing to the algorithmic decision to set the thresholds at medians of the scores derived from all evaluated variants identified in the Pan-Cancer and ExAC cohorts, which differed from the median values of variants identified in the Pan-Cancer and 1000 Genomes cohorts.

Although we excluded TCGA subset from the ExAC cohort to avoid contamination of the control with cancer patients, a large portion of the ExAC cohort was comprised of individuals with diseases that might be associated with LSD-causing mutations (e.g., schizophrenia and bipolar disorder).^12^ Furthermore, population structure adjustment was infeasible for this cohort because the individual-level genotype data were not accessible at the time we conducted this study. As shown in Supplementary Fig. 14, the mean PPV frequency varied considerably across populations in the ExAC cohort, and correlations between the PPV frequencies of different populations were relatively low for the East Asian and African populations. Therefore, results from association analyses using this cohort as control might be confounded by the population structure difference and should be interpreted with caution.

### Statistical analysis

A two-step approach was employed to examine the association between PPVs and cancer. In the first step, the Pan-Cancer and 1000 Genomes cohorts were analyzed with the SKAT-O method for the aggregate rare-variant association and Fisher’s exact tests and logistic regressions for direct comparison of mutation prevalence.^69^ The Cochran-Armitage trend test was used to evaluate the association between cancer risk and PPV load. We adjusted for population structure using principal component analysis on 10,494 tag-SNPs, as described above (Supplementary Fig. 13). In the second step, we used the ExAC cohort an independent control and performed Fisher’s exact tests to validate the preceding results. Age at diagnosis of cancer was compared using Wilcoxon rank sum tests and linear regression. We performed DEG and gene set analyses using the DESeq2 Bioconductor package and the GAGE method based on the framework of KEGG pathways, respectively.^75,77,78^

Correction for multiple testing was conducted using the FDR estimation procedure, and the tail area-based FDR (also referred to as *q*-value) was reported.^79^ All tests were two-tailed unless otherwise specified. We considered FDR<0.1 and P<0.05 (when not adjusted for multiple testing) significant. Statistical analysis was performed using R software, version 3.5.0 (R Foundation for Statistical Computing, Vienna, Austria), with packages of Bioconductor version 3.7.

## Data availability

The data that support the findings of this study are available publicly or with proper authorization. The germline and somatic (tumor) variant call sets and the RNA-Seq read count matrices derived from the PCAWG project are available for general research use under the data access policies of the ICGC and TCGA projects. In order to gain authorized access to the controlled-tier elements of the data, researchers will need to apply to the TCGA Data Access Committee via dbGAP (https://dbgap.ncbi.nlm.nih.gov/aa/wga.cgi?page=login) for the TCGA portion and to the ICGC Data Access Compliance Office (DACO) at http://icgc.org/daco for the remainder. Clinical and pathological data of individual donors and specimens are in an open tier and are accessible through the ICGC Data Portal at https://dcc.icgc.org/releases/PCAWG. For researchers who obtained authorization from the ICGC DACO, detailed instructions on data download are available at http://docs.icgc.org/pcawg/data/. Variant call sets derived from the 1000 Genomes project phase 3 and the ExAC release 1.0 are publicly available at the individual level and the population level, respectively, from the sources described in the Methods.

## Acknowledgments

The members of the PCAWG steering committee (Peter J. Campbell, Gad A. Getz, Joshua M. Stuart, Jan O. Korbel, and Lincoln D. Stein) have reviewed and approved the submission of the manuscript as a companion paper from the PCAWG Germline Cancer Genome Working Group of the ICGC/TCGA Pan-Cancer Analysis of Whole Genomes Network. This study was supported by the Korean Cancer Foundation (K20170519); Korea Health Technology R&D Project through the Korea Health Industry Development Institute, funded by the Ministry of Health & Welfare, Republic of Korea (HI14C0072 and HI14C2399); National Research Foundation through contract N-16-NM-CR01-S01; and the Program of Construction and Operation for Large-scale Science Data Center (K-16-L01-C06-S01).

## Author Contributions

J.S. and Y.K. conceptualized and designed the study. D.K. downloaded, merged, and preprocessed the variant call sets into an analyzable structure. J.S. and D.K. performed variant annotation, examination and manual curation, and final classification of the variants. J.S. performed the statistical analysis. J.S. and D.K. wrote the draft of the manuscript. M.C. and Y.K. provided critical input on data analysis. S.S.Y. held the authorized access to the PCAWG project data as a member of the Cancer Genome Project leadership at ICGC. J.O.K., as a member of the PCAWG scientific steering committee, co-chaired the germline working group and provided guidance to this study. Y.K. and S.S.Y. supervised the overall study and were responsible for the final approval of the manuscript. All authors contributed to the interpretation of results and vouch for the accuracy and integrity of the overall content.

